# Molding the Rice Methylome for Disease Resistance

**DOI:** 10.64898/2026.05.03.722557

**Authors:** Leonardo Furci, Jurriaan Ton, Hidetoshi Saze

## Abstract

In *Arabidopsis thaliana*, epigenetic changes in the DNA methylome can prime transcriptional responses to biotic and abiotic stress, resulting in enhanced resistance. Epigenetic recombinant inbred lines have further enabled the identification of trans-generationally stable epialleles controlling stress resistance without adverse effects on plant growth, highlighting their potential for crop improvement. Unfortunately, extending these approaches to crops has remained largely unsuccessful due to differences in genome architecture. The rice (*Oryza sativa*) genome consists for ∼40% of transposable elements and other epigenetically regulated repeat sequences. Therefore, perturbation of DNA methylation typically leads to severe developmental defects, sterility or lethality, which precludes the use of methylome engineering strategies for epiallele mapping and crop improvement.

Here, we exploit an inducible system to introduce for the first time widespread epigenetic variation in rice without detrimental phenotypic consequences. We combined the *A. thaliana* DNA demethylase *AtROS1* with the β-estradiol-activated *XVE* cassette (*XVE:AtROS1-YFP*) to enable transient DNA demethylation during early development. Induction of the construct in transgenic Nipponbare seedlings yielded genome-wide changes in the DNA methylome which persisted for at least one generation. Strikingly, these methylome changes did not cause developmental defects or reduced seed yield, but instead correlated with enhanced resistance against *Xanthomonas oryzae*, the causal agent of bacterial leaf blight in rice.

Our study demonstrates that controlled, transient DNA demethylation can uncouple epigenetic variation from deleterious phenotypes in rice. This approach provides a practical framework for generating epigenetic mapping populations and opens new avenues for harnessing epigenetic variation in crop improvement.

## Introduction

DNA methylation - the addition of a methyl group to the 5th carbon of cytosine residues in DNA - is a central epigenetic mark in plants. It plays fundamental roles in preserving genome stability by repressing potentially deleterious transposable elements (TEs) and fine-tuning gene expression (Mattei et al., 2022). In most plants, DNA methylation occurs at cytosines in three different sequence contexts CG, CHG and CHH (where H can be A C or T), which is established and maintained via interdependent pathways (Gwee et al., 2025). *De novo* DNA methylation is established by the RNA-dependent DNA methylation (RdDM) pathway, which utilizes 24nt small-interfering RNAs (siRNAs) produced by the plant-specific RNA polymerases Pol-IV and Pol-V to direct DNA methylase DOMAINS REARRANGED METHYLTRANSFERASE 2 (DRM2) to target sequences (Erdmann and Picard, 2020). DNA methylation at the CG context it is maintained during DNA replication across cell division by METHYLTRANSFERASE 1 (MET1), whereas at CHG sequences by CHROMOMETHYLASE 3 (CMT3) in conjunction with a self-reinforcing feedback loop with H3K9me2 histone methyltransferases SU(VAR)3–9 HOMOLOGS 4/5/6 (SUVH4/5/6). Maintenance of DNA methylation at the asymmetric CHH context is mediated by the RdDM pathway and CHROMOMETHYLASE 2 (CMT2), which requires the chromatin remodeler DEFICIENT IN DNA METHYLATION 1 (DDM1) (Gwee *et al*., 2025). To prevent excessive spread of DNA methylation and unwanted silencing of necessary genes, a family of bifunctional DNA glycosylases/lyases act as DNA demethylases by cleaving the methylated cytosine from the DNA backbone and initiating base-excision repair. Among these, REPRESSOR OF SILENCING 1 (ROS1) is mostly responsible for DNA demethylation in vegetative tissues through a combination of DNA glycosylase activity (Liu and Lang, 2020) and passive occupancy-based prevention of *de novo* DNA methylation mechanism (Deng et al., 2026).

These core mechanisms of DNA (de)methylation are conserved across angiosperms (Schmitz et al., 2019; Zhu et al., 2025), enabling dynamic and locus-specific regulation of gene expression. This regulatory flexibility underpins transcriptional reprogramming during physiological responses to fluctuating environmental conditions – a key aspect of phenotypic plasticity in sessile organisms such as plants (Miryeganeh and Saze, 2020). Indeed, whole-genome DNA methylation (the “methylome”) have been shown to change in response to both abiotic and biotic stress across several plant species. In addition, mutants in DNA methylation maintenance machinery often display modified responses to both abiotic and biotic stresses (Hannan Parker et al., 2022; Liu et al., 2022b; Lloyd and Lister, 2022). In the model plant *Arabidopsis thaliana* (Arabidopsis), ROS1 has been shown to actively remove DNA methylation at the promoter region of the immune receptor gene *RLP43* when exposed to the bacterial pathogen *Pseudomonas syringae* or bacterial elicitor *flg22* (Halter et al., 2021), or at repeats in the promoters of stress-regulatory genes *ACD6*, *ACO3*, and *GSTF14* after exposure to cold, drought or salt stress (Yang et al., 2022), altering their transcriptional responsiveness. ROS1-dependent DNA methylation has also been implicated in the priming of defense genes against herbivory (Wilkinson et al. 2023).

Stress exposure can prime plants to respond more effectively to recurring challenges, enhancing fitness with minimal metabolic costs (López Sánchez et al., 2021; van Hulten et al., 2006). This phenomenon is referred to as induced resistance in the context of biotic stress, or acquired tolerance in response to abiotic stress. Central to this process is the establishment of somatic stress memory that persists during stress-free periods, which involves histone modifications and changes in DNA methylation, as well as other physiological mechanisms(Cooper and Ton, 2022; Harris et al., 2023). In the case of biotic interactions, the primed state boosts innate immunity and pathogen resistance, which in some cases can be transmitted to isogenic progeny for one or more generations in the absence of stress. This phenomenon, termed heritable induced resistance (h-IR), has been reported across taxonomically diverse plant species, including crops (Furci et al., 2023b). Unlike animals, plants only partially reset acquired changes in DNA methylation during reproduction. As a result, stress-induced epigenetic variation can persist across generations as metastable, yet reversible, differentially methylated regions (Cao and Chen, 2024). These epialleles provide a plausible mechanistic basis for the inheritance of stress-adaptive traits, such as h-IR and abiotic stress tolerance. However, despite their potential importance, only a limited number of epialleles have been causally linked to phenotypic variation in plants; most described cases are naturally occurring and related to developmental processes rather than stress resistance (Furci et al., 2023a).

In Arabidopsis, epigenetic recombinant inbred lines (epiRILs) have been generated by crossing DNA hypomethylated mutants such as *met1* and *ddm1* with wild-type plants. Revertant wild-type progeny from selfed F_1_ hybrids maintained recombinant epialleles from the mutant parent for up to eight generations (Johannes et al., 2009; Reinders et al., 2009). These populations have enabled the mapping and linking of stably inherited epialleles with both complex developmental traits, such as flowering time or root length (Cortijo et al., 2014), as well as agronomically-relevant traits, such as pathogen resistance or salt stress tolerance, which occur without major physiological costs (Furci et al., 2019; Kooke et al., 2015; Liégard et al., 2019). These findings highlight epigenetic variation as a source of heritable phenotypic diversity that could be harnessed for crop improvement, particularly in light of emerging epigenome editing technologies (Agarwal et al., 2020; Harris et al. 2023, Yang et al., 2025).

Unfortunately, translating these approaches to crop species has proven challenging due to fundamental differences in genome organization. Compared to Arabidopsis, most crops have larger genomes with a higher proportion of transposable elements (TEs) and repetitive DNA, rendering them more sensitive to major changes in DNA methylation. For example, while ∼20% of the Arabidopsis genome consists of TEs, this proportion amounts to ∼40% in the rice (*Oryza sativa*) cultivar Nipponbare. In rice, disruption of key DNA methylation regulators, including *OsMET1-2*, *OsDDM1a/b*, *OsDRM2*, or *OsROS1*, leads to severe developmental defects such as dwarfism, sterility, and embryo lethality (Higo et al., 2012; Hu et al., 2014; Tan et al., 2016; Zhou et al., 2021). Similarly, chemical inhibition of DNA methyltransferase activity by 5-azacytidine or 5-aza-2’-deoxycytidine cause a global reduction of DNA methylation at the cost of severe developmental defects (Liu et al., 2022a; Sano et al., 1990). On the other hand, it has been suggested that larger genomes containing higher number of transposons and repetitive DNA have an elevated potential for the generation of epialleles (Catoni and Cortijo, 2018). Consistent with this potential, several naturally occurring epialleles have been identified in rice, including variants affecting developmental and stress-related traits. For example, epigenetic silencing of *DWARF1* results in a stable dwarf phenotype (Miura et al., 2009), while differential methylation at the *ACQUIRED COLD TOLERANCE 1* (*ACT1*) promoter has recently been shown to modulate heritable cold tolerance, contributing to the geographic expansion of cultivated rice (Song et al., 2025). However, the inability to experimentally induce and stably propagate methylome variation without deleterious effects has so far limited the systematic identification and exploitation of such epialleles.

Here, we address this limitation by exploiting an inducible epigenome-modifying system in rice based on the β-estradiol–responsive XVE cassette driving the expression of the Arabidopsis DNA demethylase ROS1 (Wilkinson et al. 2023; Parker et al. 2025). Transient induction of AtROS1 during early development enabled widespread DNA methylome reprogramming in Nipponbare without compromising plant growth, development, or yield. The resulting epigenetic variation was stable across somatic tissues and heritable, and was associated with enhanced resistance to *Xanthomonas oryzae*, the causal agent of bacterial leaf blight. Together, these results establish a practical framework for generating epigenetic mapping populations in rice and open new avenues for dissecting and harnessing epiallelic variation in crop improvement.

## Results and Discussion

### Introduction and characterization of XVE:ROS1-YFP vector in Nipponbare rice

The *XVE:ROS1-YFP* vector incorporates the β-estradiol-inducible *XVE* cassette (Siligato et al., 2016; Zuo et al., 2000) with the Arabidopsis DNA demethylase *ROS1* fused with the *YFP Venus* reporter tag, allowing for β-estradiol-dependent activation of transgenic *AtROS1*. This vector has been used successfully to induce global DNA methylation alterations in Arabidopsis (Hannan Parker et al., 2025; Wilkinson et al., 2023). After embryogenic calli transformation with the *XVE:ROS1-YFP* vector, a T_0_ Nipponbare heterozygote plant was propagated for two generations (T_2_) to isolate progeny homozygous for the vector insertion (XR1) and wild-type revertants as a control (NB*; Figure 1A). As calli generation and rice transformation processes have been shown to induce durable methylome changes (Hsieh et al., 2023; Stroud et al., 2013), this strategy ensures a more robust control to isolate specific DNA methylation changes arising from β-estradiol treatment in transgenic plants, and exclude lineage-specific epigenomic variations independent from *AtROS1* expression, which would arise if comparing XR1 plants with wild-type Nipponbare from a different origin.

**Figure 1:**
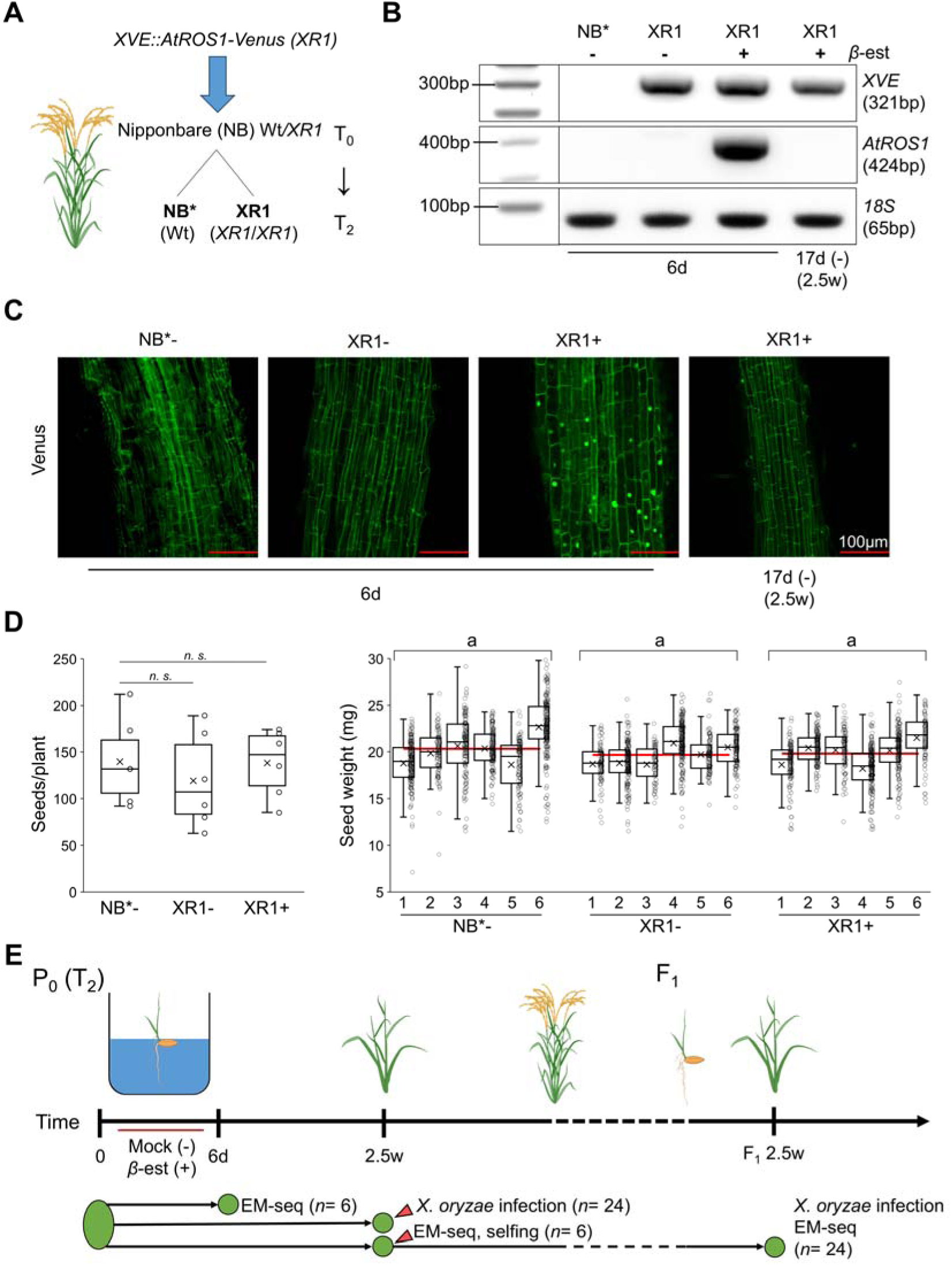
Introduction and characterization of *XVE:ROS1-YFP* vector in Nipponbare rice plants. **(A)** Diagram of transformation and selection for obtaining homozygote T_2_ transgenic lines containing the *XVE:ROS1-YFP* vector (XR1) and wild-type Nipponbare null segregant (NB*) from parental transformant line (T_0_) **(B)** Validation of β-estradiol-dependent *AtROS1* transcriptional expression in leaves of transgenic XR1 seedlings through RT-PCR, while germinated in liquid media containing 150μM β-estradiol (+) or mock (-) for 6 days (6d), and subsequent transcriptional deactivation after additional 11 days of growth in absence of β-estradiol (2.5w). **(C)** Validation of β-estradiol-dependent AtROS1 protein expression in transgenic XR1+ seedlings through tagged YFP Venus fluorescence imaging. Roots were imaged after 6d of germination in liquid media containing β-estradiol (+) or mock (-), and protein expression disappeared after additional 11 days of growth in absence of β-estradiol (2.5w; scale bar: 100µm) **(D)** Left panel: seed yields per plant for NB* and XR1 plants treated with β-estradiol (+) or mock (-) at 6d and grown until seed set. Differences in average seed number produced per plant between XR1 lines and NB* control were assessed statistically by two-tailed Student’s t-test (*n* = six plants). Right panel: individual seed weight (in mg) across β-estradiol (+) or mock (-) treated NB* and XR1 rice plants. Center line indicates median, with lower and upper quartiles below and above; small crosses indicate mean; red line indicates grand mean seed weight across the six plants. Differences in seed weights between NB*-, XR1- and XR1+ plants were assessed with a linear mixed model, with replicates as a random grouping factor using SPSS (*p*= 0.713). Post-hoc comparison were performed using Bonferroni-corrected estimated marginal means between NB*-, XR1- and XR1+ plants (small letters, *n. s*). **(E)** Diagram of experimental setup for induction and analysis of within-generation and transgenerational methylome alterations and pathogen resistance phenotyping. Green dots indicate separate batches (i.e. seedlings/plants) used in each experiment.

RT-PCR on cDNA from leaf tissue of NB* and XR1 seedlings germinated for 6 days (6d) in ½ MS media with either 150μM β-estradiol (+) or mock (-) revealed specific expression of transgenic *AtROS1* in XR1+ seedlings. Notably, after seedlings were transplanted from liquid culture and grown until 2.5 weeks of age (2.5w) in the absence of β-estradiol, *AtROS1* expression was no longer detectable in XR1+ plants (Figure 1B). Confocal laser scanning microscopy confirmed nuclear-localized fluorescence of the AtROS1-fused YFP Venus tag in roots of XR1+ plants, which has disappeared at 2.5w in the absence of β-estradiol (Figure 1C). Hence, β-estradiol-inducible *XVE:ROS1-YFP* transactivation vector is functional in Nipponbare rice plants.

Importantly, XR1 seeds germinated in ½ MS media with either 150μM β-estradiol (+) or mock (-) for 6d and then transplanted to soil could reach flowering stage and yield similar seed number and size to those of wild-type NB* plants germinated in mock-treated ½ MS media (*n*= 6, Figure 1D). This demonstrates that transient *AtROS1* expression at the seedling stage does not cause major developmental defects or affect seed yield and viability, unlike most genetic mutations and or chemical inhibitors of DNA methylation machinery.

To evaluate whether this ROS1-inducible system is capable of achieving global alterations of the rice methylome, NB* and XR1 lines (P_0_) were germinated in liquid ½ MS medium containing 150μM β-estradiol (+) or mock (-) for 6d, at which point global DNA methylation levels were examined in leaf tissues using enzymatic methyl-seq (EM-seq) (Figure 1E). This method of global DNA methylation profiling is more accurate and reliable for the detection of CG and non-CG DNA methylation than whole-genome bisulfite sequencing (Feng et al., 2020). Germinated rice seedlings were transferred to soil and grown for 2.5w in the absence of β-estradiol. To test whether ROS1-induced global changes in DNA methylation at the seedling stage can induce within-generation changes to agronomically-relevant phenotypes, a subset of 2.5w old rice plants was challenge-inoculated with *X. oryzae* pv *oryzae* (*Xoo*), the causal agent of rice bacterial leaf blight, to assess changes to resistance phenotype (Figure 1E). In parallel, a subset of 2.5w plants was analyzed for methylome changes through EM-seq (Figure 1E). Lastly, to determine heritable stability of the ROS1-induced epigenomic and phenotypic changes, F_1_ progenies of EM-sequenced NB* and XR1 plants from were grown for 2.5w in the absence of β-estradiol and tested for global methylome changes (EM-seq) and *Xoo* resistance (Figure 1E).

### Global DNA methylation changes induced by AtROS1 activation in estradiol-treated rice seedlings

To determine the effect of β-estradiol-induced *AtROS1* expression on XR1 plants the DNA methylome at 6d, genome-wide DNA methylation at single-cytosine resolution was examined in NB*-, XR1- and XR1+ seedling leaves by EM-seq analysis, using the recently released Nipponbare telomere-to-telomere genome assembly (NIP-T2T, AGIS1.0) as reference (Shang et al., 2023) (Figure 1E). At both protein coding (PC) and TE/repeat loci, XR1+ seedlings exhibited a global reduction of DNA methylation across all cytosine contexts when compared to XR1-, whereas DNA methylation of XR1-seedlings was comparable to NB* (Figure 2A, Supplemental Figure 1). As expected, at the single-cytosine level, a larger proportion of differentially methylated cytosines (DMCs) showing hypomethylation was found. The density of these hypomethylated DMCs was evenly distributed across chromosome arms and pericentromeric regions, including both PC-rich and TE-rich regions, while being absent in the centromeres of most chromosomes (Figure 2B, Supplemental Figure 2). Notably, hypermethylated DMCs were also identified, albeit at lower proportion, and their distribution showed higher density in centromeres or pericentromeric regions (Figure 2B, Supplemental Figure 2). Differentially methylated regions (DMRs), rather than individual DMCs, more likely cause stable changes in gene expression and phenotypes(Weigel and Colot, 2012). Therefore, we interrogated our EM-seq data for ROS1-induced DMRs between XR1- and XR1+ seedling leaves at 6d. As expected, a larger proportion of DNA hypomethylated DMRs was identified compared to hypermethylated ones (Figure 2C). While both hypo-and hypermethylated DMRs appeared to be more frequent in intergenic regions and TEs (Figure 2D, Venn diagrams), the unique genomic loci associated with these DMRs were enriched in PC genes relative to the total number of TEs or PC loci (Figure 2D, table). Representative loci with intergenic or genic DMRs specific to XR1+ seedlings are shown in Figure 2E.

**Figure 2:**
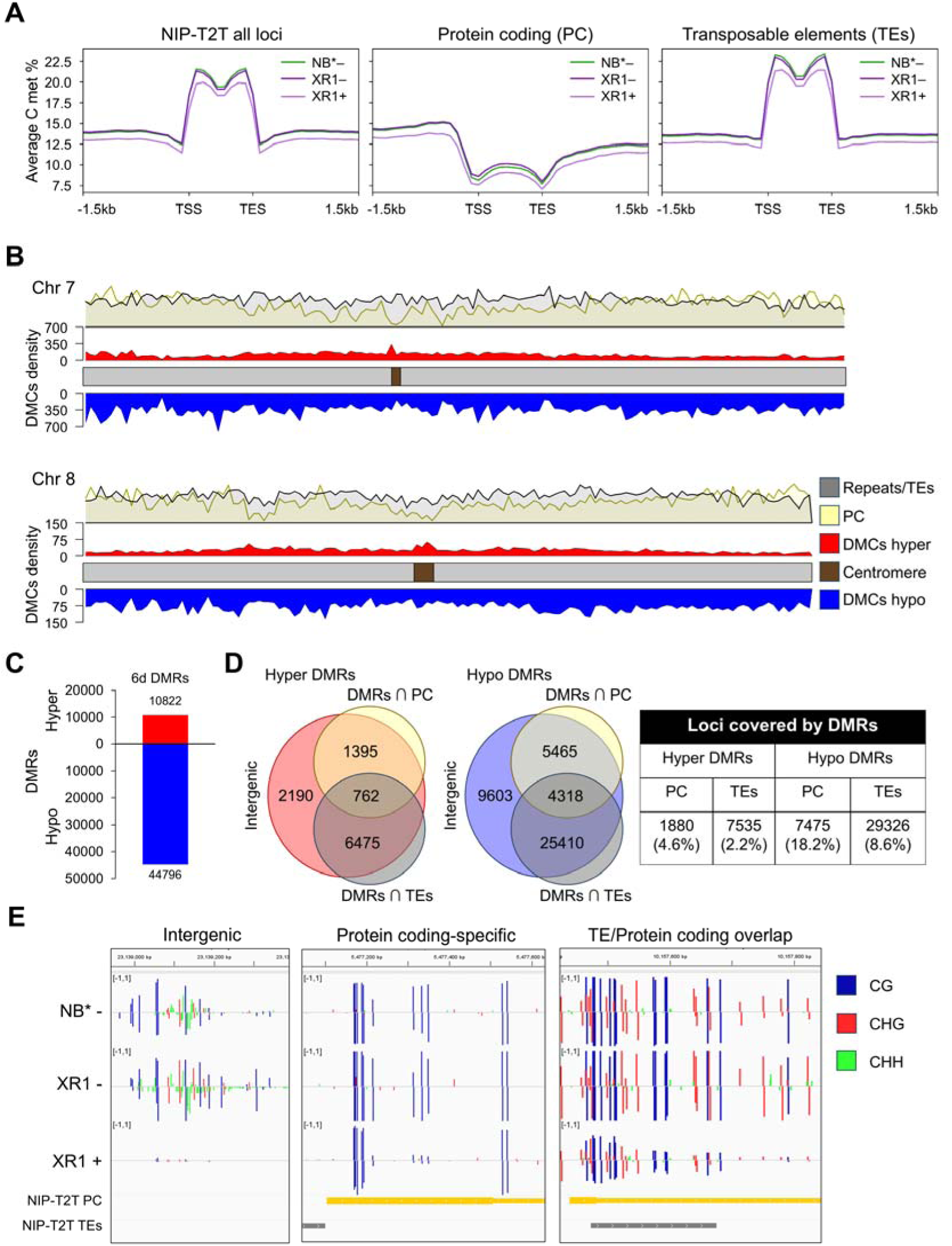
Rice methylome alterations by β-estradiol treatment in six-day old XR1 seedlings. **(A)** Metaplots displaying average DNA methylation (all C contexts) in six-days old (6d) wild-type segregant NB* or transgenic XR1 rice seedlings germinated in liquid media containing 150μM β-estradiol (+) or mock (-) across all loci of Nipponbare telomere-to-telomere assembly (NIP-T2T), protein coding genes (PC) or transposable elements (TEs)/repeat elements (AGIS1.0 annotation). **(B)** Distribution of differentially methylated cytosines (DMCs) density at representative chromosomes 7 and 8 identified in β-estradiol-treated XR1 seedlings (+) vs mock-treated ones (-), separated by hypermethylated (red) or hypomethylated (blue). Above, yellow and grey density plots show distribution frequencies of PC or TEs loci, respectively. **(C)** Number of differentially methylated regions (DMRs) in 6d XR1+ vs XR1-seedlings, separated by hypermethylated (red) or hypomethylated (blue) **(D)** Venn diagrams: overlap (∩) between unique hypo- or hypermethylated DMRs in six-days old XR1+ vs XR1-seedlings with PC or TE loci (AGIS1.0 annotation); Table: number of unique genetic loci covered by one or more DMRs (percentage in parenthesis represents the proportion over the total number of respective loci in the annotation). **(E)** Browser view examples (IGV) of DMRs overlapping intergenic regions (left panels), PC gene (RAP ID Os09g0118666, middle panel) or both PC and TE (Locus ID LOC_Os02g17600, right panel). Blue bars: methylation % at cytosines in CG context, red bars: methylation % at cytosines in CHG contexts, green bars: methylation % at cytosines in CHH contexts. Positive and negative values represent methylation % of cytosines on the positive or negative strand, respectively.

The rice *Osmet1-2* mutant predominantly affects CG methylation (Hu *et al*., 2014), whereas *ddm1* mutations in both rice and Arabidopsis mostly affect DNA methylation in pericentromeric TEs (Tan *et al*., 2016; Zemach et al., 2013). Our results highlight the advantages of using the *XVE:ROS1-YFP* vector in achieving global changes in DNA methylation across all Cs and all genetic loci, when compared to molecular genetic approaches using knock-outs of DNA methylation maintenance genes.

Rice plants treated with 5-aza-2’-deoxycytidine showed hypermethylated DMRs which correlated with an increased abundance of 24-nt siRNA (Liu *et al*., 2022a). Additionally, a recent study using the *XVE:ROS1-YFP* vector in Arabidopsis showed an increase of pericentromeric DMCs following β-estradiol treatment, which was accompanied by transcriptional activation of *CLASSY3* in leaves (Hannan Parker *et al*., 2025).

CLASSY3 recruits Pol-IV and initiates the RdDM pathway, and its coding gene is normally expressed in reproductive tissues. These findings strongly suggest that the (mostly pericentromeric) DNA hypermethylation found in XR1+ seedlings following β-estradiol-dependent AtROS1 induction is caused by *de novo* DNA methylation through RdDM pathway, most likely initiated by siRNA-generating loci de-repressed by the removal of DNA methylation.

### Within-generation maintenance of altered DNA methylation yields resistance to X. oryzae

To investigate the within-generation metastability of DNA methylation alterations in XR1 plants at the seedling stage, a subset of six XR1- and XR1+ plants treated at 6d were grown to 2.5w in the absence of β-estradiol, and leaf tissue was collected to perform methylome analysis through EM-seq (Figure 1E). In XR1+ plants, DMCs were still present across all chromosomes. However, a comparison of DMCs distribution between XR1+ and XR1-plants at 6d versus 2.5w revealed an overall reduction of hypomethylated DMCs, across all chromosomes. (Figure 3A, Supplemental Figure 3).

**Figure 3:**
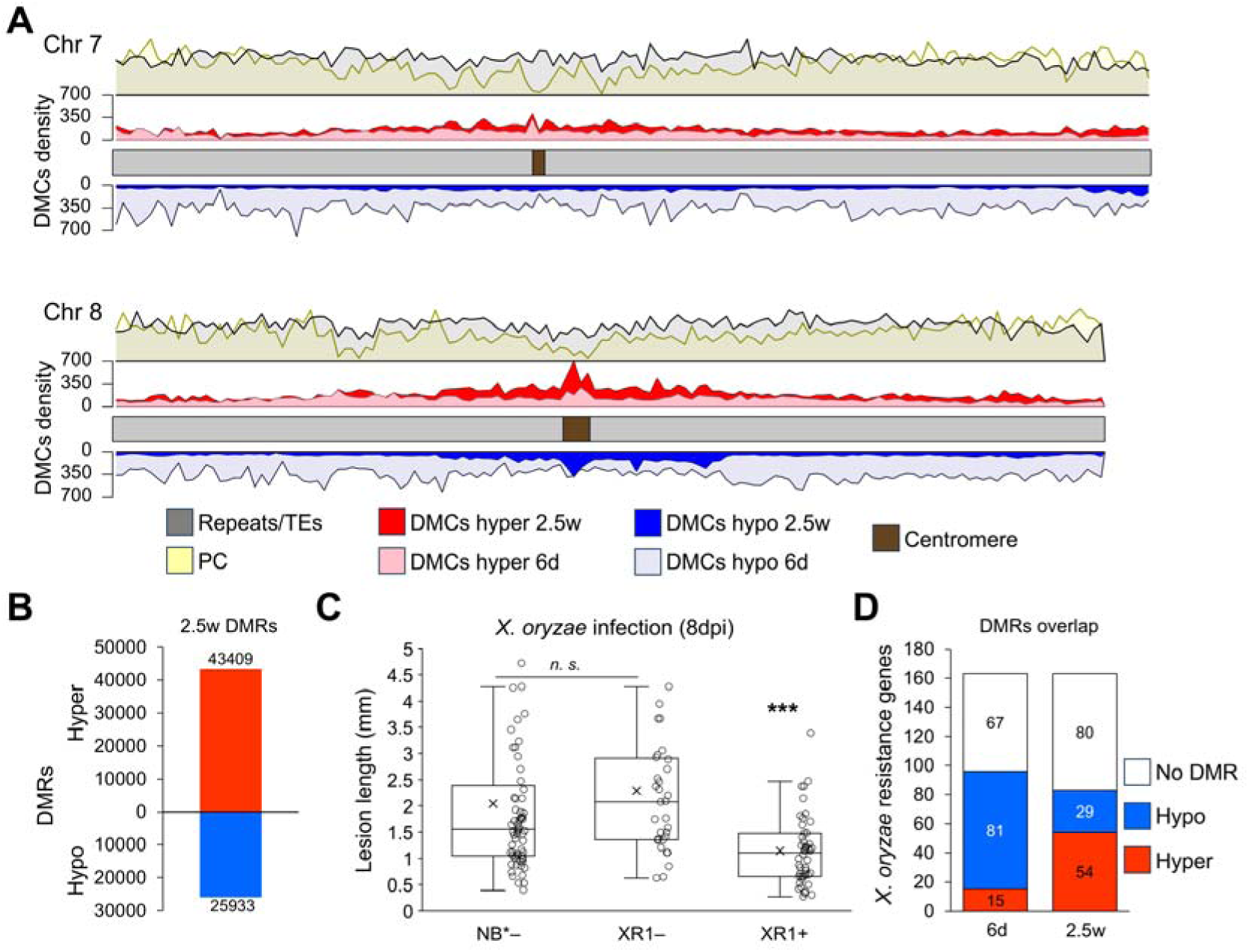
Methylome alterations in 2.5w old plants treated with β-estradiol at seedling stage and *X. oryzae* resistance phenotyping. **(A)** Distribution of methylated cytosines (DMCs) density at representative chromosomes 7 and 8 identified between 2.5 weeks-old (2.5w) XR1 plants treated with 150μM β-estradiol (+) at six days-old (6d) vs mock-treated ones (-), separated by hypermethylated (red) or hypomethylated (blue). Lighter red and blue colors represent DMCs distribution in 6d seedlings (from Figure 2A) for comparison. Above, yellow and grey density plots show distribution frequencies of PC or TE loci, respectively. **(B)** Number of differentially methylated regions (DMRs) identified in 2.5w XR1+ plants vs XR1-ones, separated by hypermethylated (red) or hypomethylated (blue). **(C)** Basal resistance phenotype of 2.5w NB* and XR1 plants against *X. oryzae* infection. Displayed are lesion lengths (in mm) in rice leaves caused by the pathogen eight days post-inoculation (dpi) though the leaf clipping method. Center line indicates median, with lower and upper quartiles below and above; small crosses indicate mean. Statistically significant differences in lesion length between XR1 lines and NB* control were assessed by two-tailed Student’s t-test (*n* =30+ leaves from 15+ plants, *** *p-*value <0.001). **(D)** Overlap between DMRs identified in 6d and 2.5w XR1+ plants with 163 genes causally linked with *X. oryzae* basal resistance genes reported in (Tonnessen *et al*., 2019).

Interestingly, chromosomes 4, 5 and 8 retained higher numbers of hypomethylated DMCs at the centromere (Figure 3A, Supplemental Figure 3), consistent with the fact that these three chromosomes possess abnormal/aberrant centromeres depleted of CentO satellite repeats but enriched with TEs (Shang *et al*., 2023). On the other hand, hypermethylated DMCs increased in centromeres, pericentromeric regions and chromosome arms (Figure 3A, Supplemental Figure 3), likely due to the *de novo* establishment of RdDM-dependent DNA methylation. As a result, a larger number of hypermethylated DMRs was identified in XR1+ plants versus XR1- at 2.5w compared to hypomethylated DMRs (Figure 3B).

To assess whether within-generation metastable methylome alterations affect agronomically-relevant traits, 2.5w old NB* and XR1 plants germinated in either 150µm β-estradiol (+) or mock (-) were infected with *Xoo* through leaf clipping (Methods; Figure 1E). At eight days post-inoculation (dpi), lesion length caused by *Xoo* infection was significantly lower in 2.5w old XR1+ plants (Figure 3C), indicating that DNA methylome alterations at the seedling stage are sufficient to induce resistance to later *Xoo* infection. A previous study reported a cluster of 163 co-expressed genes after *Xoo* infection, which causally contribute to basal resistance against this pathogen (Tonnessen et al., 2019). More than half of those genes (83/163) were found to contain one or more DMRs in their gene body or promoter (1000 bp window upstream of the transcription start site) in 2.5w XR1+ plants (Figure 3D), suggesting a direct link between ROS1-induced changes in DNA methylation and the disease resistance phenotype.

### Transgenerational metastable DMRs of rice lines maintaining transgenerational resistance to X. oryzae map to defense genes

The six rice plants per genotype/condition whose methylome was sequenced at 2.5w were grown further to seed maturation stage to obtain the next generation (F_1_) seeds, to estimate the transgenerational metastability of parental methylome alterations when germinated in either β-estradiol (P_0_+) or mock (P_0_-) (Figure 1E).

EM-seq methylome analysis of individual F_1_ XR1 P_0_+ lines (named XR1-1 to XR1-6) versus XR1 F_1_ P_0_-revealed a large number of both DNA hypo- and hypermethylated DMRs in all XR1 P_0_+ lines. Furthermore, by comparing the genomic location and DNA methylation direction of DMRs in each of the F_1_ XR1 P_0_+ lines against the DMRs of their corresponding XR1+ parental lines, we could identify between 5.1% and 17.6% of DMRs present in both P_0_ and F_1_ (Figure 4A, grey boxes). For all lines, both DNA hypo-and hypermethylated DMRs overlapping between P_0_ and F_1_ plants were significantly higher than expected by chance after 10,000 random shuffling permutations (Figure 4A, white asterisks). We therefore conclude that these regions are stably transmitted across meiosis into the next generation.

**Figure 4:**
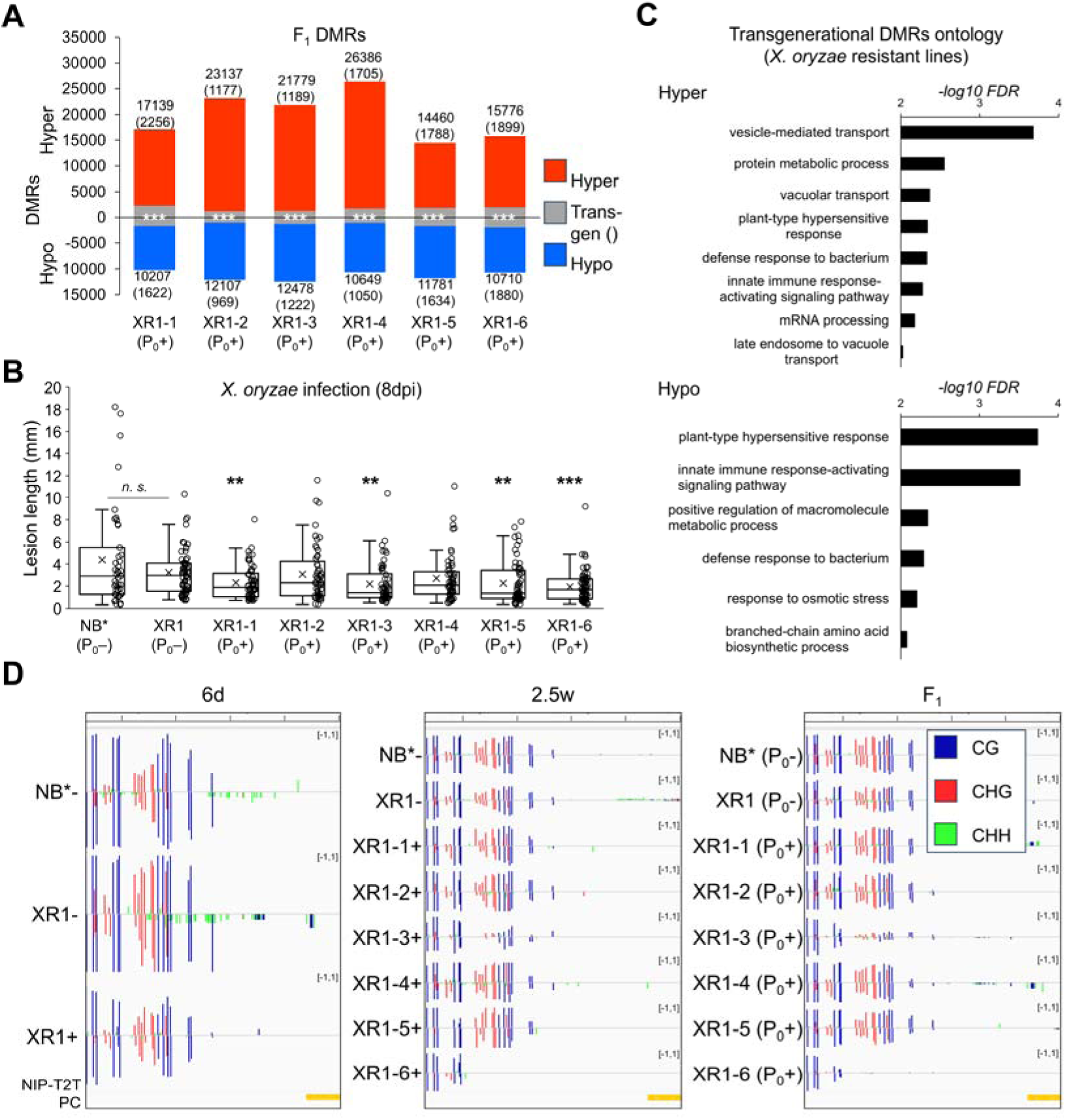
Transgenerational stability of methylome alterations and *X. oryzae* resistance in the progeny generation of parental XR1 lines treated with β-estradiol at the seedling stage. **(A)** Number of differentially methylated regions (DMRs) identified in 2.5 weeks-old (2.5w) progeny XR1 plants (F_1_) derived from parental lines treated with β-estradiol at their seedling stage (P_0_+, XR1-1 to XR1-6) vs parental lines treated with mock (P_0_-), separated by hypermethylated (red) or hypomethylated (blue). Gray boxes indicate the number of hypo- or hypermethylated DMRs overlapping between P_0_ and F_1_ (transgenerational DMRs). For all lines, the number of transgenerational DMRs have been found to be significantly higher than expected than occurring by chance after random shuffling of the genomic regions (10000 permutations, *** *p-*value <0.001). **(B)** Basal resistance phenotype of F_1_ 2.5w NB* and XR1 plants against *X. oryzae* infection. Displayed are lesion lengths (in mm) in rice leaves caused by the pathogen eight days post-inoculation (dpi) though the leaf clipping method. Center line indicates median, with lower and upper quartiles below and above; small crosses indicate mean. Statistically significant differences in lesion length between XR1 lines and NB* control were assessed in pairwise comparisons by two-tailed Student’s t-test (*n* =40+ leaves from 20+ plants, ** *p-*value <0.01, *** *p-*value <0.001). **(C)** Ontology terms enrichment (biological processes) for genes overlapped by transgenerational hypo- and hypermethylated DMRs unique to *X. oryzae*-resistant lines XR1-1, XR1-4, XR1-4 and XR1-6. **(D)** Browser view example (IGV) of transgenerational DMRs maintained as DNA hypomethylated epiallele in line XR1-3 and XR1-6 across one generation, located in the promoter of *USP37* (RAP ID Os10g0463300, yellow bar below represents the first exon). Blue bars: methylation % at cytosines in CG context, red bars: methylation % at cytosines in CHG contexts, green bars: methylation % at cytosines in CHH contexts. Positive and negative values represent methylation % of cytosines on the positive or negative strand, respectively.

To assess whether the heritable DMRs contribute to disease resistance, the same F_1_ plants were inoculated with *Xoo* (Figure 1E). At eight dpi, four XR1 P_0_+ lines displayed a statistically significant reduction in lesion lengths compared to NB* or XR1 P_0_-plants (Figure 4B, asterisks), indicating heritable transmission of the ROS1-induced resistance against *Xoo*. On the other hand, two XR1 P_0_+ lines showed lesion length comparable to NB* or XR1 P_0_-plants (Figure 4B), indicating a reversion of P_0_ *Xoo* resistance to the susceptibility levels of wild-type NB* or P_0_-XR1, which is a plausible outcome for an epigenetically controlled quantitative trait. Indeed, the heritable DMRs specific to the resistant lines (XR1-1, XR1-3, XR1-5 and XR1-6) overlapped with genes that are statistically enriched with gene ontology terms related to plant immunity and pathogen resistance (Figure 5C), whereas the heritable DMRs specific to the susceptible lines (XR1-2 and XR1-4) failed to show any significant enrichment for biological function ontologies. An example of a DNA hypomethylated DMR inherited as an epiallele is shown in Figure 4D, located in the promoter of the universal stress protein A (UspA) domain-containing protein coding gene *USP37* (RAP ID Os10g0463300). Together, these results reveal a clear link between the heritable epialleles and the heritable induced resistance phenotype against *Xoo*.

In conclusion, our study has demonstrated that the *XVE:ROS1-YFP* vector can be used to epigenetically improve crop resistance without accompanying costs on physiological growth or seed yield. Some of the DMRs caused by transient *AtROS1* induction at seedling stage were stably inherited to the next generation and correlated with enhanced resistance to rice leaf blight. We therefore propose the *XVE:ROS1-YFP* construct can be exploited as a novel molecular tool to generate epiRIL populations and fine-map the epigenetic bases of agronomically important complex traits in crops.

## Methods

### Rice growth conditions and β-estradiol treatment

Rice plants were grown in Biotron LPH-411PFDT-S growth cabinets (Nihonika, Japan) under ∼600μmol/s/m^-1^ illumination. For vegetative growth, plants were grown at long-day condition, with 16h light at 28°C and 70% relative humidity (RH), followed by 8h of darkness at 24°C and 70% RH. After 45 days of vegetative growth, plants were switched to short-day conditions to induce flowering, with 12h light at 28°C and 70% RH, followed by 12h of darkness at 24°C and 70% RH until seed production.

Dehusked and surface-sterilizes rice seeds (70% EtOH, 1 min) were germinated in 50ml of sterile ½ strength liquid MS media (Sigma-Aldrich, + 0.01% MES, pH 5.8) supplemented with β-estradiol or mock. Powdered β-estradiol (TCI chemicals) was dissolved to 100mM stock concentration using DMSO (Fujifilm Wako chemicals). For seed germination, 75μl of 100mM β-estradiol stock were added to 50ml of ½ strength MS media, whereas 75μl of pure DMSO was added as mock.

### RT-PCR

Total RNA was extracted from 6d rice seedling leaves using the Maxwell 16 LEV Plant RNA kit (Promega) according to the manufacturer instructions. One μg of total RNA was then converted to cDNA using PrimeScript II 1st strand cDNA Synthesis Kit (Takara). All primers used are listed in Supplemental Table 1.

### Fluorescent imaging

Root samples from 6d seedlings were fixed in 4% paraformaldehyde (in PBS) for 30 min and then suspended in ClearSee solution (Kurihara et al., 2015) for six days at room temperature for tissue clearing. Fixed and cleared roots were imaged on Nikon A1 confocal microscope.

### EM-seq and data analysis

High-purity genomic DNA was extracted using Illustra™ Nucleon™ PhytoPure™ kit (Cytiva), then sheared to ∼500bp fragments through sonication using Covaris M220. Enzymatic conversion of methylated cytosines and library preparation were performed using NEBNext® Enzymatic Methyl-seq Kit and NEBNext® Multiplex Oligos for Enzymatic Methyl-seq (New England Biolabs) according to the provided instructions, using sodium hydroxide for DNA denaturation and performing libraries size selection (470-520bp) for enrichment of larger inserts. Libraries were sequenced on Illumina NovaSeq X Plus using 2x 150bp paired-end configuration.

Reads were adaptor-trimmed and paired using Trimmomatic (Bolger et al., 2014) with settings LEADING:10 TRAILING:5 SLIDINGWINDOW:4:20 MINLEN:33. Reads alignment to NIP-T2T genome (Shang *et al*., 2023) and DNA methylation calling was done with Bismark 0.23.0 (Krueger and Andrews, 2011), using de-deduplication and all-C methylation extractor.

Across all experimental conditions, DMCs were called by assessing significant difference in methylation at each cytosine position between XR1+ and XR1-plants through Chi-squared test (≥5 reads per position, ≥20% DNA methylation proportion variation and *p-*value <0.01). Chromosomal distribution of DMCs was visualized using the R package “karyoploteR” (bin size 200000bp) (Gel and Serra, 2017).

DMRs were identified with DMRcaller (Catoni et al., 2018) using Fisher’s exact test with *bin* method using the following settings: bin size 50bp, minimum reads per cytosine 4, minimum cytosine count 2, false discovery rate-adjusted *p-*value (Benjamini-Hochberg method) <0.05, minimum proportion DNA methylation difference CG ≥40%, CHG ≥40% CHH ≥20%.

For both DMCs and DMRs, all significant position/regions identified between XR1-and XR1+ samples were cross-referenced against differentially methylated position/regions identified in a comparison between NB* samples versus Nipponbare rice of different genetic origin, to exclude potential lineage-specific or stochastic DMCs or DMRs. The NIP-T2T AGIS1.0 protein coding genes and TEs/repeats annotations were used for downstream analysis of DMRs overlaps.

Significant overlaps in transgenerational DMRs between P_0_ and F_1_ lines were estimated using the R package “regioneR” (Gel et al., 2016) using the *randomizeRegions* function to check against random distributions (10000 permutations).

Significant Gene ontology enrichment for Biological Processes terms was performed with PANTHER (v19.0, Benjamini–Hochberg fdr-adjusted *p*-value <0.01)(Thomas et al., 2022).

### X. oryzae resistance assays

*Xanthomonas oryzae* pv. *oryzae* strain T7174 (*Xoo*) was obtained from National Agriculture and Food Research Organization (NARO) Genebank, Japan (MAFF ID: 311018). *Xoo* leaf clipping inoculation was carried out as explained in (Ke et al., 2017). Briefly, frozen *Xoo* stock was grown on solid peptone sucrose agar (PSA) media (for one liter: 10g peptone, 1g L-glutamic acid, 10g sucrose, pH 7) at 28°C for three days until a biofilm was formed. Then, *Xoo* biofilm was scraped and grown in 50ml of liquid PSA at 28°C for two additional days. Before inoculation, *Xoo* was re-suspended in sterile 10mM MgCl_2_ to a final OD_600_ of 0.5. The two largest leaves from each 2.5w old plants were then clipped 2-3cm from the tip with scissors dipped in the *Xoo-*MgCl_2_ suspension. At eight dpi, lesion lengths from *Xoo* infection were quantified from high-resolution pictures of infected rice leaves using ImageJ imaging software (Schneider et al., 2012).

## Data availability

Illumina EM-seq data (adaptor-trimmed *.fastq* files) generated in this study have been deposited in EMB-EBI European Nucleotide Archive database under accession code PRJEB111188.

## Acknowledgments

This work has been supported by MEXT Grant-in-Aid for Transformative Research Areas (A) JP20H05913 to H.S., JSPS Grant-in-Aid for Early-Career Scientists 23K13957 to L.F., by JSPS Grant-in-Aid for JSPS Fellows 23KF0288 to L. F., and a BBSRC-IPA grant (BB/W015250/1) and ERC grant (PantMemo; 101199639) to JT.

We are grateful for the help and support provided by the Sequencing Section (SQC) of Core Facilities at Okinawa Institute of Science and Technology Graduate University for Illumina sequencing of EM-seq samples.

We are grateful for the help and support provided by Dr. Paolo Barzaghi (Paolone) from the Imaging Section of Core Facilities at Okinawa Institute of Science and Technology for fluorescent imaging of YFP Venus reporter in rice seedling roots.

---

**Supplemental Figure 1:**
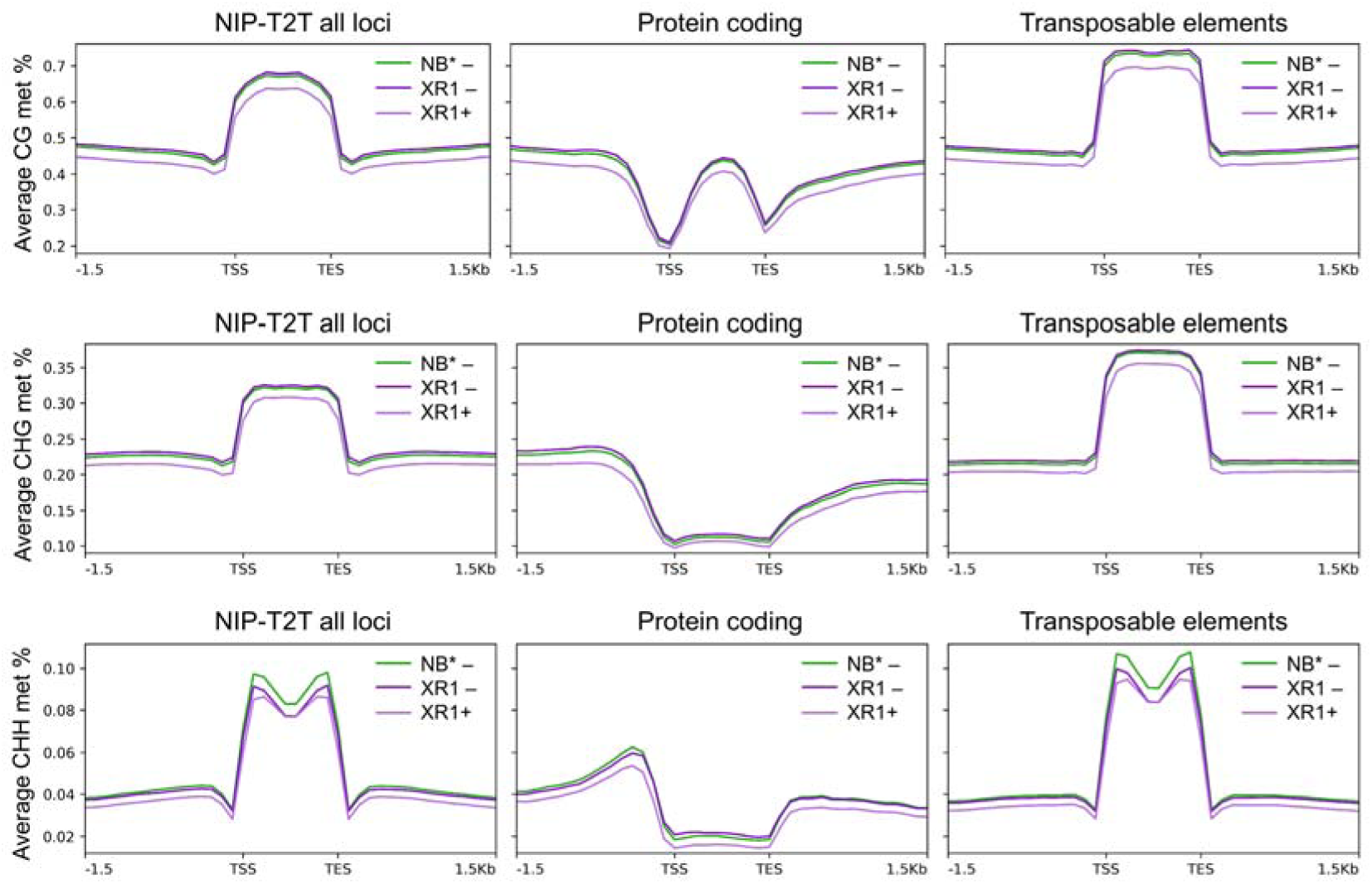
Metaplots displaying average DNA methylation separated by CG, CHG or CHH contexts in six-days old (6d) wild-type segregant NB* or transgenic XR1 rice seedlings germinated in liquid media containing 150μM β-estradiol (+) or mock (-) across all loci of Nipponbare telomere-to-telomere assembly (NIP-T2T), protein coding genes (PC) or transposable elements (TEs)/repeat elements (AGIS1.0 annotation).

**Supplemental Figure 2:**
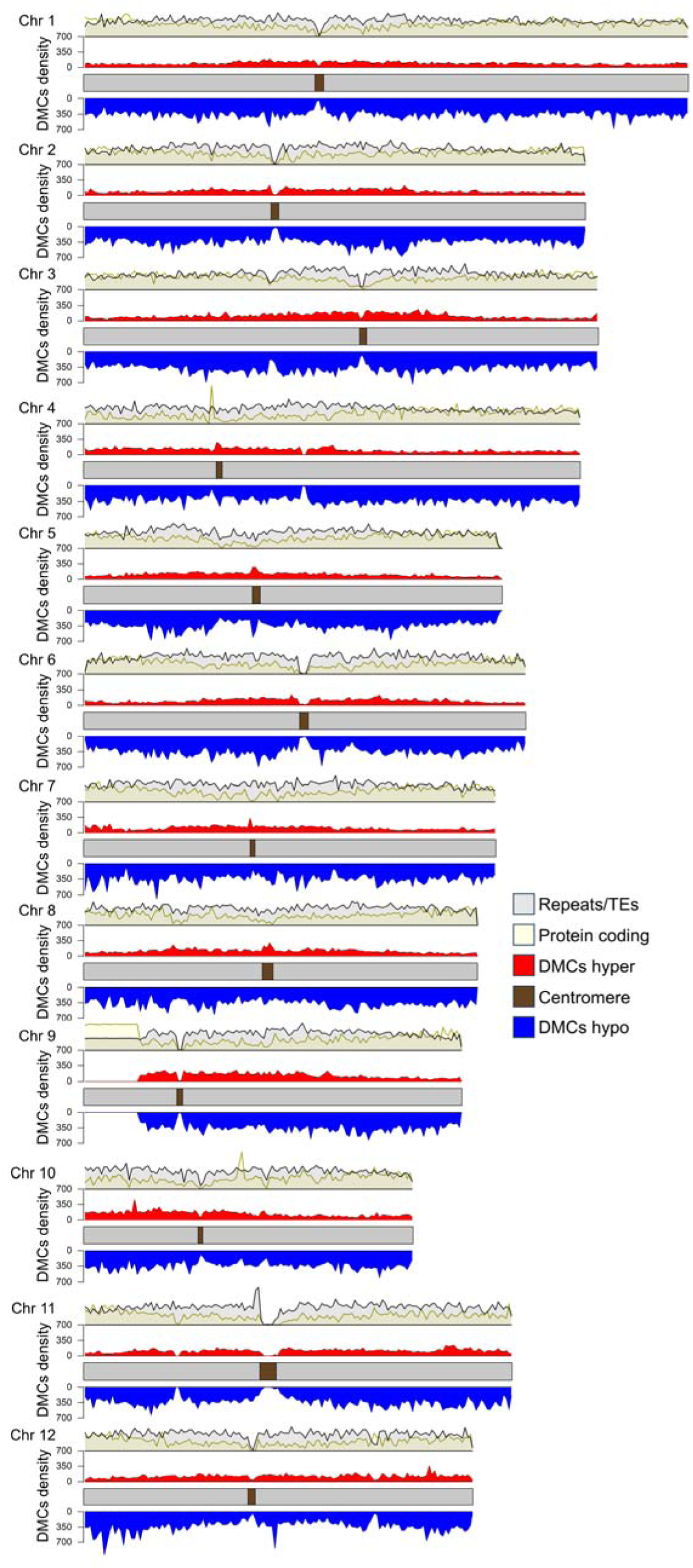
Distribution of differentially methylated cytosines (DMCs) density at all Nipponbare chromosomes identified in β-estradiol-treated XR1 seedlings (+) vs mock-treated ones (-), separated by hypermethylated (red) or hypomethylated (blue). Above, yellow and grey density plots show distribution frequencies of PC or TEs loci, respectively.

**Supplemental Figure 3:**
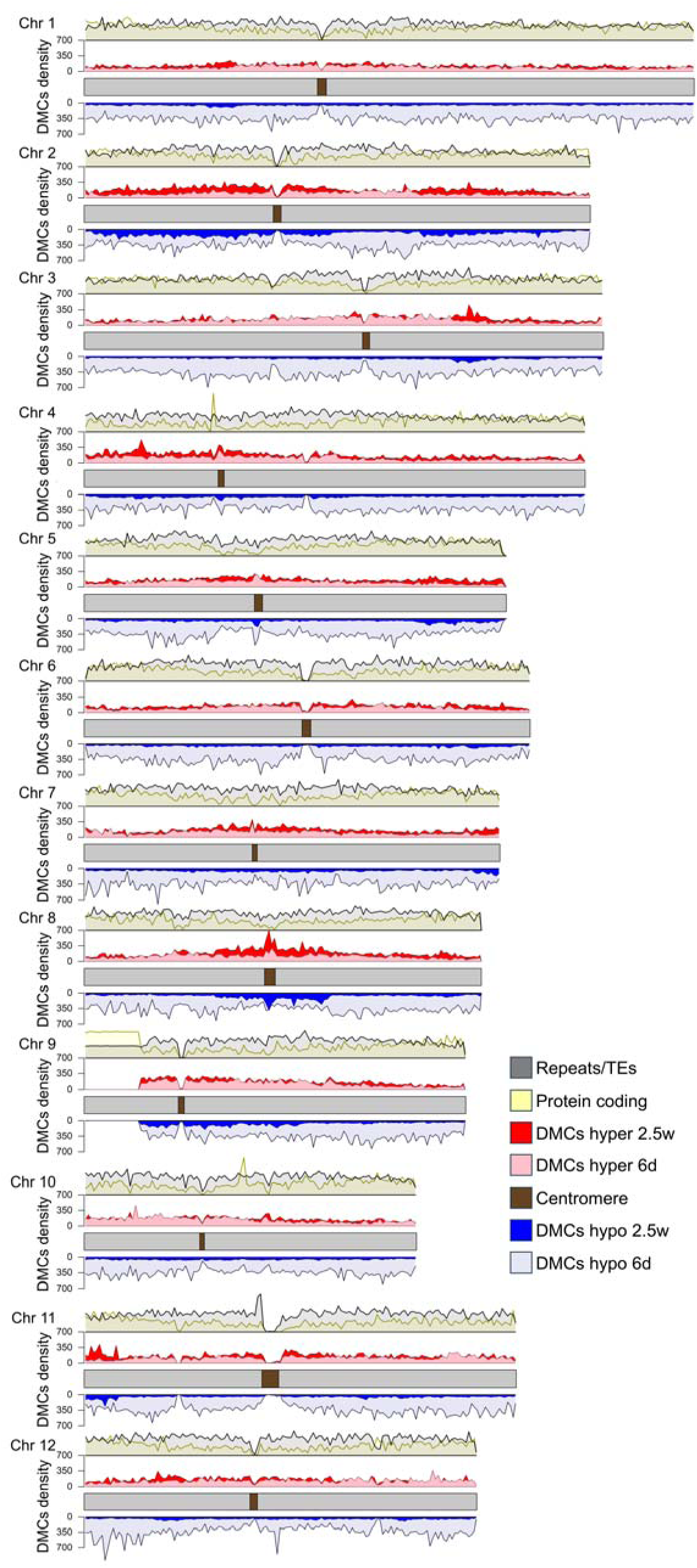
Distribution of methylated cytosines (DMCs) density at all Nipponbare chromosomes identified between 2.5 weeks-old (2.5w) XR1 plants treated with 150μM β-estradiol (+) at six days-old (6d) vs mock-treated ones (-), separated by hypermethylated (red) or hypomethylated (blue). Lighter red and blue colors represent DMCs distribution in 6d seedlings (from Supplemental Figure 2) for comparison. Above, yellow and grey density plots show distribution frequencies of PC or TE loci, respectively.

## References

Bolger, A.M., Lohse, M., and Usadel, B. (2014). Trimmomatic: a flexible trimmer for Illumina sequence data. Bioinformatics 30:2114–2120. 10.1093/bioinformatics/btu170.

Cao, S., and Chen, Z.J. (2024). Transgenerational epigenetic inheritance during plant evolution and breeding. Trends in Plant Science 29:1203–1223. 10.1016/j.tplants.2024.04.007.

Catoni, M., and Cortijo, S. (2018). Chapter Four - EpiRILs: Lessons From Arabidopsis. In Advances in Botanical Research, M. Mirouze and E. Bucher and P. Gallusci, eds. (Academic Press: pp. 87–116. 10.1016/bs.abr.2018.08.002.

Catoni, M., Tsang, J.M., Greco, A.P., and Zabet, Nicolae R. (2018). DMRcaller: a versatile R/Bioconductor package for detection and visualization of differentially methylated regions in CpG and non-CpG contexts. Nucleic Acids Research 46:e114–e114. 10.1093/nar/gky602.

Cooper, A., and Ton, J. (2022). Immune priming in plants: from the onset to transgenerational maintenance. Essays Biochem 66:635–646. 10.1042/ebc20210082.

Cortijo, S., Wardenaar, R., Colomé-Tatché, M., Gilly, A., Etcheverry, M., Labadie, K., Caillieux, E., Hospital, F., Aury, J.-M., Wincker, P., et al. (2014). Mapping the Epigenetic Basis of Complex Traits. Science 343:1145–1148. doi:10.1126/science.1248127.

Deng, L., Zhu, G., Zhong, W., Jia, Z., Zhou, Q., Zhang, M., Si, Z., Zhang, Q., Liang, Y., Du, X., et al. (2026). Occupancy-based mechanism is the chief mode of ROS1 function in preventing DNA hypermethylation. Nature Plants 10.1038/s41477-026-02258-z.

Erdmann, R.M., and Picard, C.L. (2020). RNA-directed DNA Methylation. PLoS Genet 16:e1009034. 10.1371/journal.pgen.1009034.

Feng, S., Zhong, Z., Wang, M., and Jacobsen, S.E. (2020). Efficient and accurate determination of genome-wide DNA methylation patterns in Arabidopsis thaliana with enzymatic methyl sequencing. Epigenetics Chromatin 13:42. 10.1186/s13072-020-00361-9.

Furci, L., Berthelier, J., Juez, O., Miryeganeh, M., and Saze, H. (2023a). Chapter 15 - Plant Epigenomics. In Handbook of Epigenetics (Third Edition), T.O. Tollefsbol, ed. (Academic Press: pp. 263–286. 10.1016/B978-0-323-91909-8.00007-4.

Furci, L., Pascual-Pardo, D., Tirot, L., Zhang, P., Hannan Parker, A., and Ton, J. (2023b). Heritable induced resistance in Arabidopsis thaliana: Tips and tools to improve effect size and reproducibility. Plant Direct 7:e523. 10.1002/pld3.523.

Furci, L., Jain, R., Stassen, J., Berkowitz, O., Whelan, J., Roquis, D., Baillet, V., Colot, V., Johannes, F., and Ton, J. (2019). Identification and characterisation of hypomethylated DNA loci controlling quantitative resistance in Arabidopsis. eLife 8:e40655. 10.7554/eLife.40655.

Gel, B., and Serra, E. (2017). karyoploteR: an R/Bioconductor package to plot customizable genomes displaying arbitrary data. Bioinformatics 33:3088–3090. 10.1093/bioinformatics/btx346.

Gel, B., Díez-Villanueva, A., Serra, E., Buschbeck, M., Peinado, M.A., and Malinverni, R. (2016). regioneR: an R/Bioconductor package for the association analysis of genomic regions based on permutation tests. Bioinformatics 32:289–291. 10.1093/bioinformatics/btv562.

Gwee, J., Tian, W., Qian, S., and Zhong, X. (2025). DNA methylation dynamics: patterns, regulation, and function. Current Opinion in Plant Biology 88:102787. 10.1016/j.pbi.2025.102787.

Halter, T., Wang, J., Amesefe, D., Lastrucci, E., Charvin, M., Singla Rastogi, M., and Navarro, L. (2021). The Arabidopsis active demethylase ROS1 cis-regulates defence genes by erasing DNA methylation at promoter-regulatory regions. eLife 10:e62994. 10.7554/eLife.62994.

Hannan Parker, A., Wilkinson, S.W., and Ton, J. (2022). Epigenetics: a catalyst of plant immunity against pathogens. New Phytologist 233:66–83. 10.1111/nph.17699.

Hannan Parker, A., Zhang, P., Man, K.Y., Tirot, L., Rolfe, S.A., Smith, L.M., Wilkinson, S.W., and Ton, J. (2025). From recall to reset: the role of DNA (de)methylation in modulating plant immune memory. bioRxiv:2025.2008.2019.671022. 10.1101/2025.08.19.671022.

Harris, C.J., Amtmann, A., and Ton, J. (2023). Epigenetic processes in plant stress priming: Open questions and new approaches. Current Opinion in Plant Biology 75:102432. 10.1016/j.pbi.2023.102432.

Higo, H., Tahir, M., Takashima, K., Miura, A., Watanabe, K., Tagiri, A., Ugaki, M., Ishikawa, R., Eiguchi, M., Kurata, N., et al. (2012). DDM1 (decrease in DNA methylation) genes in rice (Oryza sativa). Mol Genet Genomics 287:785–792. 10.1007/s00438-012-0717-5.

Hsieh, J.-W.A., Chang, P., Kuang, L.-Y., Hsing, Y.-I.C., and Chen, P.-Y. (2023). Rice transformation treatments leave specific epigenome changes beyond tissue culture. Plant Physiology 193:1297–1312. 10.1093/plphys/kiad382.

Hu, L., Li, N., Xu, C., Zhong, S., Lin, X., Yang, J., Zhou, T., Yuliang, A., Wu, Y., Chen, Y.R., et al. (2014). Mutation of a major CG methylase in rice causes genome-wide hypomethylation, dysregulated genome expression, and seedling lethality. Proc Natl Acad Sci U S A 111:10642–10647. 10.1073/pnas.1410761111.

Johannes, F., Porcher, E., Teixeira, F.K., Saliba-Colombani, V., Simon, M., Agier, N., Bulski, A., Albuisson, J., Heredia, F., Audigier, P., et al. (2009). Assessing the Impact of Transgenerational Epigenetic Variation on Complex Traits. PLOS Genetics 5:e1000530. 10.1371/journal.pgen.1000530.

Ke, Y., Hui, S., and Yuan, M. (2017). Xanthomonas oryzae pv. oryzae Inoculation and Growth Rate on Rice by Leaf Clipping Method. Bio Protoc 7:e2568. 10.21769/BioProtoc.2568.

Kooke, R., Johannes, F., Wardenaar, R., Becker, F., Etcheverry, M., Colot, V., Vreugdenhil, D., and Keurentjes, J.J.B. (2015). Epigenetic Basis of Morphological Variation and Phenotypic Plasticity in Arabidopsis thaliana. The Plant Cell 27:337–348. 10.1105/tpc.114.133025.

Krueger, F., and Andrews, S.R. (2011). Bismark: a flexible aligner and methylation caller for Bisulfite-Seq applications. Bioinformatics 27:1571–1572. 10.1093/bioinformatics/btr167.

Kurihara, D., Mizuta, Y., Sato, Y., and Higashiyama, T. (2015). ClearSee: a rapid optical clearing reagent for whole-plant fluorescence imaging. Development 142:4168–4179. 10.1242/dev.127613.

Liégard, B., Baillet, V., Etcheverry, M., Joseph, E., Lariagon, C., Lemoine, J., Evrard, A., Colot, V., Gravot, A., Manzanares-Dauleux, M.J., et al. (2019). Quantitative resistance to clubroot infection mediated by transgenerational epigenetic variation in Arabidopsis. New Phytol 222:468–479. 10.1111/nph.15579.

Liu, R., and Lang, Z. (2020). The mechanism and function of active DNA demethylation in plants. Journal of Integrative Plant Biology 62:148–159. 10.1111/jipb.12879.

Liu, S., Bao, Y., Deng, H., Liu, G., Han, Y., Wu, Y., Zhang, T., and Chen, C. (2022a). The Methylation Inhibitor 5-Aza-2′-Deoxycytidine Induces Genome-Wide Hypomethylation in Rice. Rice 15:35. 10.1186/s12284-022-00580-6.

Liu, Y., Wang, J., Liu, B., and Xu, Z.-Y. (2022b). Dynamic regulation of DNA methylation and histone modifications in response to abiotic stresses in plants. Journal of Integrative Plant Biology 64:2252–2274. 10.1111/jipb.13368.

Lloyd, J.P.B., and Lister, R. (2022). Epigenome plasticity in plants. Nature Reviews Genetics 23:55–68. 10.1038/s41576-021-00407-y.

López Sánchez, A., Pascual-Pardo, D., Furci, L., Roberts, M.R., and Ton, J. (2021). Costs and Benefits of Transgenerational Induced Resistance in Arabidopsis. Frontiers in Plant Science Volume 12–202110.3389/fpls.2021.644999.

Mattei, A.L., Bailly, N., and Meissner, A. (2022). DNA methylation: a historical perspective. Trends in Genetics 38:676–707. 10.1016/j.tig.2022.03.010.

Miryeganeh, M., and Saze, H. (2020). Epigenetic inheritance and plant evolution. Population Ecology 62:17–27. 10.1002/1438-390X.12018.

Reinders, J., Wulff, B.B., Mirouze, M., Marí-Ordóñez, A., Dapp, M., Rozhon, W., Bucher, E., Theiler, G., and Paszkowski, J. (2009). Compromised stability of DNA methylation and transposon immobilization in mosaic Arabidopsis epigenomes. Genes Dev 23:939–950. 10.1101/gad.524609.

Sano, H., Kamada, I., Youssefian, S., Katsumi, M., and Wabiko, H. (1990). A single treatment of rice seedlings with 5-azacytidine induces heritable dwarfism and undermethylation of genomic DNA. Molecular and General Genetics MGG 220:441–447. 10.1007/BF00391751.

Schmitz, R.J., Lewis, Z.A., and Goll, M.G. (2019). DNA Methylation: Shared and Divergent Features across Eukaryotes. Trends in Genetics 35:818–827. 10.1016/j.tig.2019.07.007.

Schneider, C.A., Rasband, W.S., and Eliceiri, K.W. (2012). NIH Image to ImageJ: 25 years of image analysis. Nature Methods 9:671–675. 10.1038/nmeth.2089.

Shang, L., He, W., Wang, T., Yang, Y., Xu, Q., Zhao, X., Yang, L., Zhang, H., Li, X., Lv, Y., et al. (2023). A complete assembly of the rice Nipponbare reference genome. Molecular Plant 16:1232–1236. 10.1016/j.molp.2023.08.003.

Siligato, R., Wang, X., Yadav, S.R., Lehesranta, S., Ma, G., Ursache, R., Sevilem, I., Zhang, J., Gorte, M., Prasad, K., et al. (2016). MultiSite Gateway-Compatible Cell Type-Specific Gene-Inducible System for Plants. Plant Physiol 170:627–641. 10.1104/pp.15.01246.

Stroud, H., Ding, B., Simon, S.A., Feng, S., Bellizzi, M., Pellegrini, M., Wang, G.L., Meyers, B.C., and Jacobsen, S.E. (2013). Plants regenerated from tissue culture contain stable epigenome changes in rice. Elife 2:e00354. 10.7554/eLife.00354.

Tan, F., Zhou, C., Zhou, Q., Zhou, S., Yang, W., Zhao, Y., Li, G., and Zhou, D.X. (2016). Analysis of Chromatin Regulators Reveals Specific Features of Rice DNA Methylation Pathways. Plant Physiol 171:2041–2054. 10.1104/pp.16.00393.

Thomas, P.D., Ebert, D., Muruganujan, A., Mushayahama, T., Albou, L.-P., and Mi, H. (2022). PANTHER: Making genome-scale phylogenetics accessible to all. Protein Science 31:8–22. 10.1002/pro.4218.

Tonnessen, B.W., Bossa-Castro, A.M., Mauleon, R., Alexandrov, N., and Leach, J.E. (2019). Shared cis-regulatory architecture identified across defense response genes is associated with broad-spectrum quantitative resistance in rice. Scientific Reports 9:1536. 10.1038/s41598-018-38195-x.

van Hulten, M., Pelser, M., van Loon, L.C., Pieterse, C.M., and Ton, J. (2006). Costs and benefits of priming for defense in Arabidopsis. Proc Natl Acad Sci U S A 103:5602–5607. 10.1073/pnas.0510213103.

Weigel, D., and Colot, V. (2012). Epialleles in plant evolution. Genome Biology 13:249. 10.1186/gb-2012-13-10-249.

Wilkinson, S.W., Hannan Parker, A., Muench, A., Wilson, R.S., Hooshmand, K., Henderson, M.A., Moffat, E.K., Rocha, P.S.C.F., Hipperson, H., Stassen, J.H.M., et al. (2023). Long-lasting memory of jasmonic acid-dependent immunity requires DNA demethylation and ARGONAUTE1. Nature Plants 9:81–95. 10.1038/s41477-022-01313-9.

Yang, L., Lang, C., Wu, Y., Meng, D., Yang, T., Li, D., Jin, T., and Zhou, X. (2022). ROS1-mediated decrease in DNA methylation and increase in expression of defense genes and stress response genes in Arabidopsis thaliana due to abiotic stresses. BMC Plant Biology 22:104. 10.1186/s12870-022-03473-4.

Zemach, A., Kim, M.Y., Hsieh, P.-H., Coleman-Derr, D., Eshed-Williams, L., Thao, K., Harmer, Stacey L., and Zilberman, D. (2013). The Arabidopsis Nucleosome Remodeler DDM1 Allows DNA Methyltransferases to Access H1-Containing Heterochromatin. Cell 153:193–205. 10.1016/j.cell.2013.02.033.

Zhou, S., Li, X., Liu, Q., Zhao, Y., Jiang, W., Wu, A., and Zhou, D.-X. (2021). DNA demethylases remodel DNA methylation in rice gametes and zygote and are required for reproduction. Molecular Plant 14:1569–1583. 10.1016/j.molp.2021.06.006.

Zhu, R., Xue, Y., and Qian, W. (2025). Molecular mechanisms and biological functions of active DNA demethylation in plants. Epigenetics Chromatin 18:41. 10.1186/s13072-025-00605-6.

Zuo, J., Niu, Q.W., and Chua, N.H. (2000). Technical advance: An estrogen receptor-based transactivator XVE mediates highly inducible gene expression in transgenic plants. Plant J 24:265–273. 10.1046/j.1365-313x.2000.00868.x.

